# Regulation of T cell receptor signaling by protein acyltransferase DHHC21

**DOI:** 10.1101/2020.02.10.942078

**Authors:** Ying Fan, Bieerkehazhi Shayahati, Ritika Tewari, Darren Boehning, Askar M. Akimzhanov

**Affiliations:** Department of Biochemistry and Molecular Biology, University of Texas-McGovern Medical School, 6431 Fannin Street, Houston, Texas 77030; Cooper Medical School of Rowan University, 401 Broadway, Camden, NJ 08103.

**Keywords:** T-cell, T helper cells, DHHC21, acyltransferase, protein acylation, protein palmitoylation, cell signaling, signal transduction

## Abstract

S-acylation – reversible post-translational lipidation of cysteine residues – is emerging as an important regulatory mechanism in T cell signaling. Dynamic S-acylation is critical for protein recruitment into the T cell receptor complex and initiation of the subsequent signaling cascade. However, the enzymatic control of protein S-acylation in T cells remains poorly understood. Here, we report a previously uncharacterized role of DHHC21, a member of the mammalian family of DHHC protein acyltransferases, in regulation of the T cell receptor pathway. We found that loss of DHHC21 prevented S-acylation of key T cell signaling proteins, resulting in disruption of the early signaling events and suppressed expression of T cell activation markers. Furthermore, downregulation of DHHC21 prevented activation and differentiation of naïve T cells into effector subtypes. Together, our study provides the first direct evidence that DHHC protein acyltransferases can play an essential role in regulation of T cell-mediated immunity.

---

T cell activation is a complex process critical for proliferation, regulation, and proper acquisition of T cell effector function. It is initiated upon ligation of the T cell receptor (TCR) by the peptide/major histocompatibility complex presented by the antigen-presenting cell. Engagement of the TCR triggers a cascade of signaling events that ultimately results in specific functional outcomes modulated by co-stimulatory molecules and cytokines (1, 2). The Src family tyrosine kinase Lck is among the first signaling molecules activated in response to TCR stimulation. Lck is rapidly recruited to the TCR where it phosphorylates TCR-associated CD3 and ζ-chain immunoreceptor tyrosine-based activation motifs (2, 3). Phosphorylation of these motifs creates a docking site for another regulatory kinase, ZAP70, resulting in its binding to the TCR and subsequent activation. Activated ZAP70 proceeds to phosphorylate its substrates, including a critically important adaptor protein LAT (linker for activation of T cells) (4). Phosphorylated LAT in turn recruits a number of signaling proteins into a multiprotein complex (“signalosome”), promoting a signaling cascade leading to T cell activation and differentiation(2, 5, 6).

Several critical regulators of the TCR signaling pathway were found to be S-acylated. Protein S-acylation, also referred to as palmitoylation, is a reversible post-translational lipidation of cysteine residues with long-chain fatty acids via a labile thioester bond. Lck was among the first mammalian proteins that were identified as S-acylated (7). Although S-acylated cysteine residues of Lck were found to be dispensable for its catalytic activity, lipidation-deficient mutants of Lck were unable to propagate TCR signaling and failed to reconstitute T cell activation in Lck-deficient cell lines (8–10). Furthermore, our previous study demonstrated that initiation of Fas-mediated apoptosis in T cells relies on rapid agonist-induced changes in Lck S-acylation levels and inhibition of Lck lipidation rendered T cells to be resistant to Fas-mediated apoptosis (11). Similar to Lck, S-acylation of LAT was found to be critically important for the initiation of downstream signaling events. A selective defect in LAT S-acylation was associated with a state of T cell functional unresponsiveness known as T cell anergy (12–14). A number of other proteins with pivotal roles in T cell activation were also discovered to be S-acylated and this post-translational modification was shown to be important for their function (15–17).

S-acylation of the mammalian proteins is catalyzed by a large family of more than 23 protein acyl transferases (PATs) with a common DHHC (Asp-His-His-Cys) motif within the catalytic center and two to six transmembrane domains (18–21). The majority of DHHC proteins are localized to ER and Golgi membranes, with a small number present at the plasma membrane (22, 23). There is evidence linking DHHC PATs to cancer, neurological and cardiovascular disorders (24–26). However, physiological roles and the protein substrate specificity of the majority of these enzymes remain enigmatic and despite the established importance of protein S-acylation for the T cell function, PATs involved in regulation of the TCR signaling pathway have not been identified.

Co-transfection experiments performed in HEK293T cells showed that DHHC21 had apparent acyltransferase activity toward Lck (27). Furthermore, we have previously reported that DHHC21 is required for activation of Lck in response to stimulation of Fas receptor in T cells (11). These observations identify DHHC21 as a strong candidate for the protein acyltransferase mediating activation of the TCR signaling pathway through S-acylation of Lck and, possibly, other T cell signaling proteins. In this study, we aimed to perform a comprehensive analysis of the DHHC21 function in T cell activation by investigating the effects of DHHC21 downregulation on S-acylation of T cell regulatory proteins, initiation of the TCR signaling pathway and expression of the T cell activation markers. Furthermore, since the T cell effector function relies on efficient TCR signal transduction, we examined whether DHHC21 is required for differentiation of naïve CD4^+^ T cells into T helper (Th) Th1 and Th2 subtypes. Overall, our study provides the first direct evidence for the regulatory role of DHHC protein acyltransferases in TCR signaling.

## Results

Given the critical importance of S-acylation for Lck signaling function, we hypothesized that DHHC protein acyltransferases are involved in initiation of the early TCR signaling events. To test this hypothesis, we treated murine EL-4 T cells with cross-linked anti-CD3 and anti-CD28 antibodies to activate the TCR signaling pathway. We then assessed lipidation of Lck cysteine residues using acyl resin-assisted capture (Acyl-RAC), a method based on selective cleavage of the thioester bond by neutral hydroxylamine followed by capture of the liberated thiols by thiopropyl sepharose (28). We found that stimulation of the TCR resulted in rapid but transient increase of S-acylation of Src-family kinases Lck and Fyn, peaking at approximately 5 minutes (Fig. 1A). Furthermore, we found that engagement of the TCR also triggered similar changes in S-acylation of phospholipase C-γ1 (PLC-γ1) (Fig. 1A), a protein that has not been previously reported to be S-acylated, suggesting that other important regulators of T cell signaling can be regulated through TCR-mediated changes in protein S-acylation levels.

**Figure 1.**
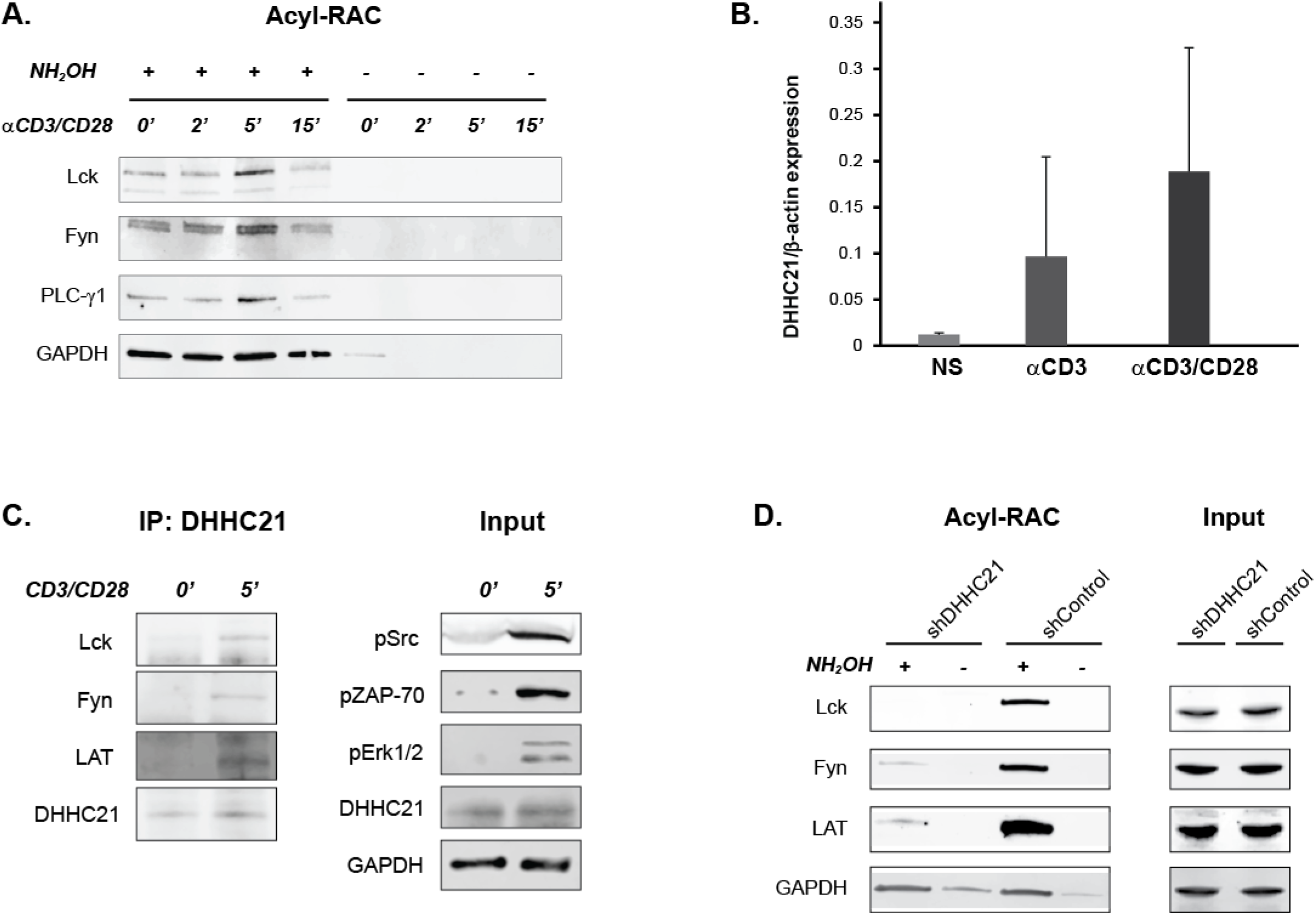
DHHC21 mediates S-acylation of T cell signaling proteins. **(A)** TCR-dependent S-acylation of signaling proteins. EL-4 T cells were stimulated with cross-linked anti-CD3/CD28 antibodies for indicated times and protein S-acylation was determined using Acyl-RAC assay. Hydroxylamine minus (NH2OH–) lanes are negative controls and represent background. GAPDH, a known S-acylated protein, was used as a loading control. **(B)** Quantitative RT-PCR analysis of DHHC21 expression in naïve murine CD4^+^ cells, unstimulated (NS) or stimulated with plate-bound anti-CD3 and anti-CD3/CD28 antibodies. DHHC21 expression shown normalized to β-actin. **(C)** Acyl-RAC analysis of protein S-acylation in EL-4 T cells transduced with DHHC21-directed or control shRNA lentivirus. **(D)** TCR-dependent interaction of DHHC21 with signaling proteins. DHHC21 was immunoprecipitated from EL-4 T cells stimulated with cross-linked anti-CD3/CD28 and binding of proteins to DHHC21 was detected by immunoblotting. Activation of the TCR pathway was confirmed by probing input lines using phospho-Src, phospho-ZAP70, and phospho-ERK1/2 antibodies.

We next sought to identify a protein acyltransferase responsible for S-acylation of Lck. Earlier, we found that protein acyltransferase DHHC21 is essential for activation of the Lck-dependent apoptotic signaling pathway in T cells (11). Our gene expression analysis revealed that DHHC21 mRNA is indeed upregulated in murine lymphoid tissues (Fig. S1) and DHHC21 expression in primary CD4^+^ T cells increased in response to TCR stimulation (Fig. 1B). Furthermore, co-expression of DHHC21 and Lck in HEK293T cells was shown to increase S-acylation of Lck, suggesting that Lck could be a protein substrate of DHHC21 (27). However, it remained unclear whether endogenously expressed DHHC21 is required for S-acylation of Lck.

To establish the protein substrate specificity of DHHC21, we first examined whether direct interaction between endogenously expressed DHHC21 and Lck in T cells can be identified by the co-immunoprecipitation assay. As shown in Fig. 1C, binding of Lck to DHHC21 cannot be confidently detected in resting EL-4 T cells. However, stimulation of the TCR with anti-CD3/CD28 antibodies significantly enhanced co-immunoprecipitation of DHHC21 with Lck and two other proximal TCR signaling proteins known to be S-acylated, Fyn and LAT (Fig. 1C).

To further confirm acyltransferase activity of DHHC21 toward proximal T cell signaling proteins, we transduced murine EL-4 T cells with lentivirus carrying either DHHC21-targeting or control shRNA vectors. All tested shRNA vectors resulted in significant (>50%) reduction of DHHC21 mRNA and protein levels in comparison to the non-targeting control (Fig S2). The effect of DHHC21 downregulation on protein lipidation was assessed using Acyl-RAC. We found that while DHHC21 knockdown did not affect total protein expression levels, it resulted in dramatic decrease of Lck, Fyn and LAT S-acylation suggesting that DHHC21 acts as a primary acyltransferase for these proteins (Fig. 1D).

S-acylation of Lck, Fyn and LAT has been shown to be essential for their signaling function (7–11, 29), suggesting that DHHC21 can regulate activation of the TCR pathway through lipidation of these proteins. To investigate whether DHHC21 is required for initiation of the early signaling events in the TCR pathway, we transduced EL-4 T cells with DHHC21-targeting shRNA and stimulated the TCR with anti-CD3/CD28 antibodies. We found that shRNA-mediated downregulation of DHHC21 resulted in decreased phosphorylation of several key signaling proteins, both on basal levels and in response to TCR stimulation (Fig. 2A). Furthermore, we observed that knockdown of DHHC21 prevented TCR-induced mobilization of calcium, another important mediator of T cell signaling (Fig. 2B).

**Figure 2.**
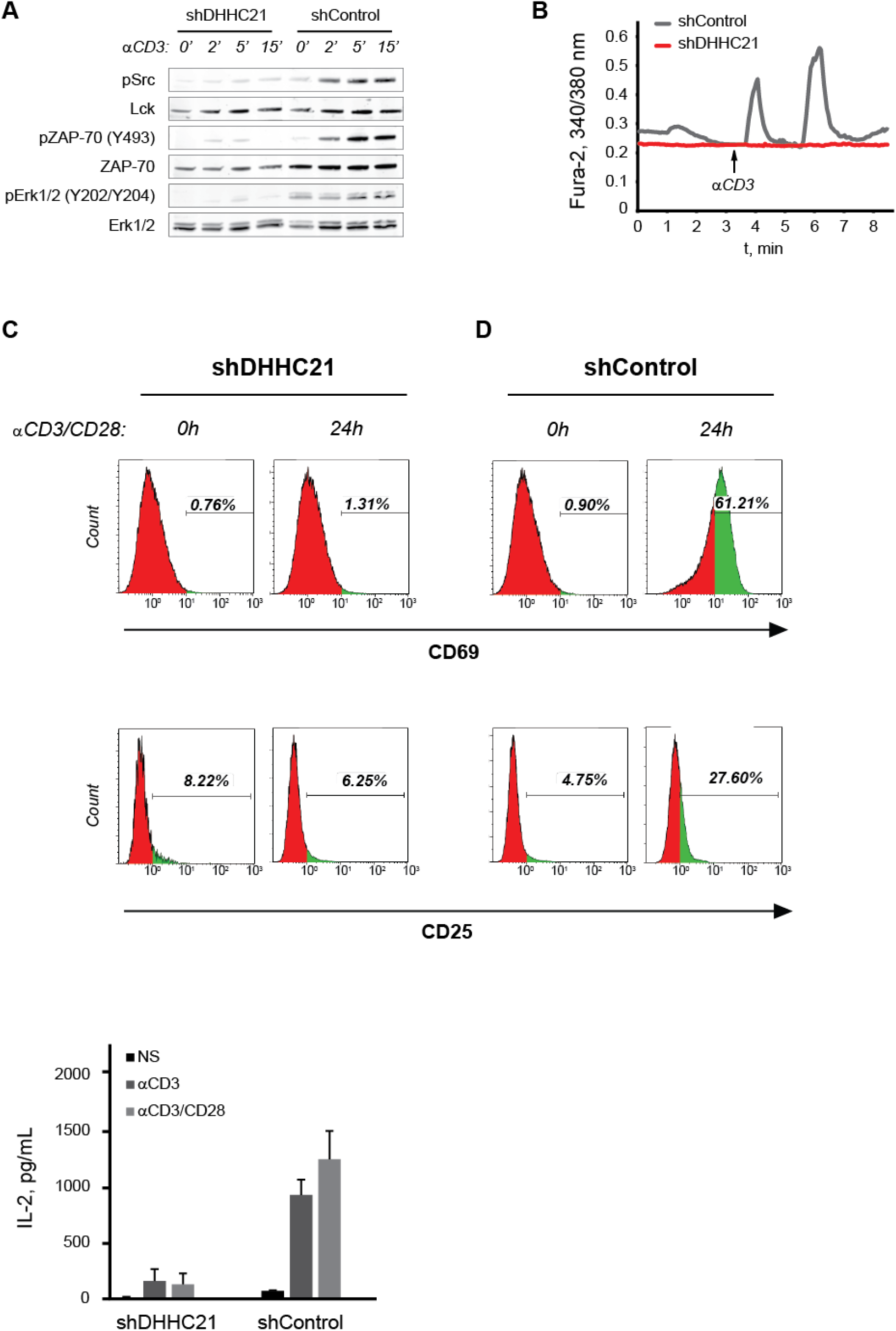
DHHC21 is required for activation of the TCR signaling pathway. EL-4 T cells were transduced with DHHC21-directed (shDHHC21) or nonspecific control (shControl) shRNA lentivirus. **(A)** Activation of TCR signaling proteins in EL-4 T cells in response to TCR stimulation with cross-linked anti-CD3/CD28 antibodies. **(B)** Fura-2 calcium imaging of EL-4 T cells stimulated with cross-linked anti-CD3/CD28 antibodies. Shown are representative single-cell responses. **(C)** Surface expression of CD69 and CD25 by EL-4 T cells stimulated with plate-bound anti-CD3/CD28 antibodies for 24 h. **(D)** IL-2 production by EL-4 T cells in response to TCR stimulation measured by ELISA.

Cytoplasmic calcium influx orchestrates early signaling events triggered by TCR engagement, ultimately leading to T cell activation and clonal expansion (30). This process is marked by a sharp increase in interleukin-2 (IL-2) production and upregulation of T cell surface receptors CD25 and CD69. To determine the functional consequences of DHHC21 downregulation on T cell activation, shRNA-transduced EL-4 T cells were stimulated with plate-bound CD3/CD28 antibodies for 24 hours and assayed for expression of the activation markers. We found that knockdown of DHHC21 completely abolished secretion of IL-2 in response to TCR treatment (Fig. 2C). Similarly, DHHC21 deficiency resulted in loss of CD25 and CD69 expression in stimulated cells (Fig. 2D) suggesting that DHHC21 is essential for T cell activation.

TCR-dependent T cell activation is a critical part of the adaptive immune response to pathogen exposure. It mediates T cell proliferation and differentiation into effector subtypes (31, 32). In particular, stimulation of the TCR triggers differentiation of naïve CD4^+^ T cells into functionally distinct subsets of effector T helper (Th) cells, such as Th1 and Th2 lineages. To test whether DHHC21 is required for development of primary T cells into Th effectors, we knocked down DHHC21 expression in isolated naïve murine CD4^+^ T cells and incubated them under Th1 and Th2 polarizing conditions. After four days, cells were re-stimulated with plate-bound CD3/CD28 antibodies and expression of Th1 and Th2-specific cytokines was measured using ELISA. We found that CD4^+^ T cells transduced with a control shRNA vector demonstrated increased production of interferon-γ (IFNγ) and IL-2 under Th1 conditions and increased levels of IL-4, IL-13 and IL-6 under Th2 conditions indicating successful polarization into the corresponding lineages (Fig. 3). Knockdown of DHHC21, however, resulted in poor production of both Th1 and Th2-specific cytokines suggesting that DHHC21 is necessary for development of the effector T helper cells.

**Figure 3.**
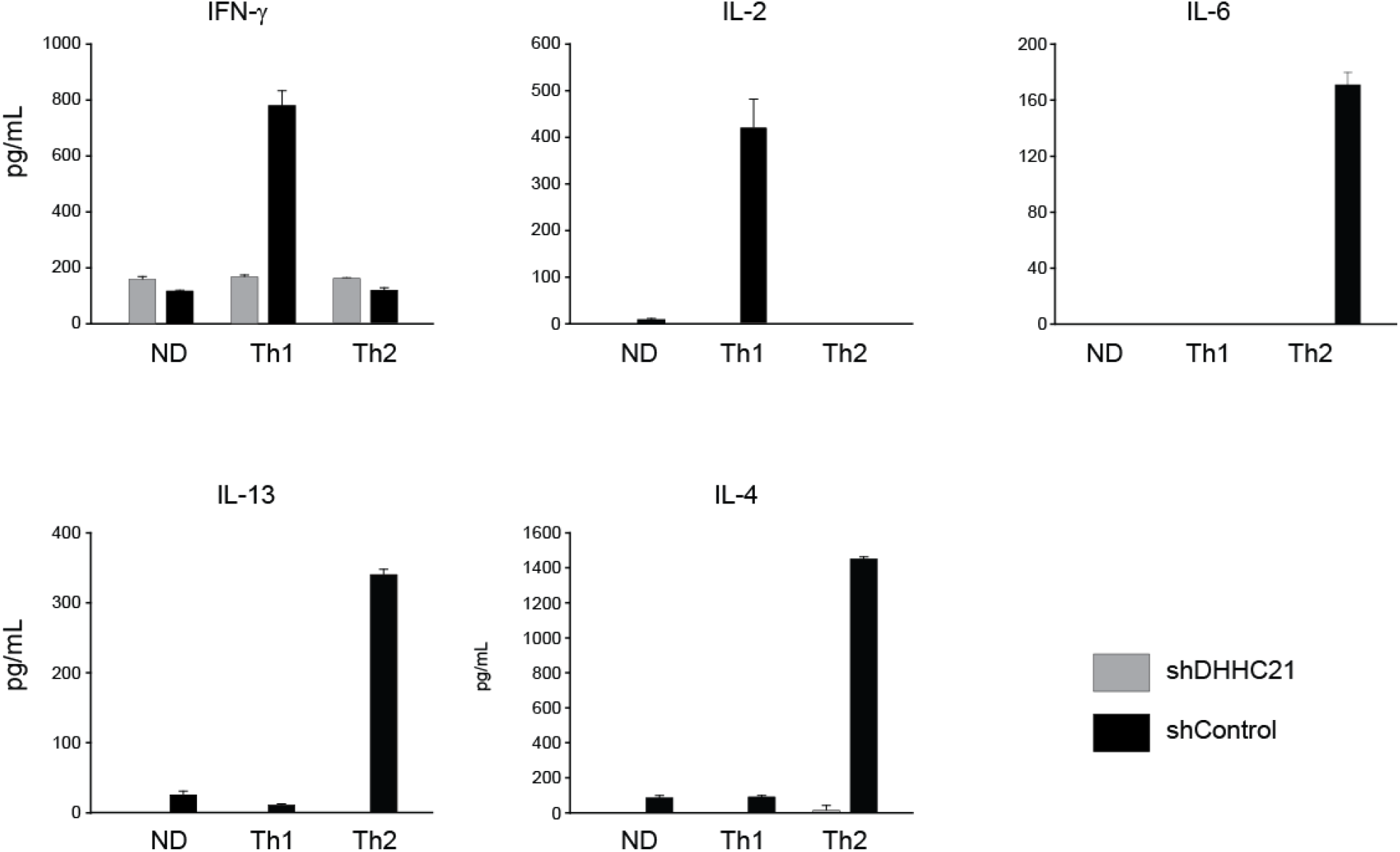
DHHC21 regulates activation differentiation of primary CD4^+^ T cells. MACS-purified naïve primary CD4^+^ T cells were transduced with DHHC21-directed (shDHHC21) or nonspecific control (shControl) ShRNA lentivirus and treated with plate-bound anti-CD3/CD28 antibodies and incubated under neutral (ND) or polarizing (Th1, Th2) conditions for 4 days. Dofferentiated CD4^+^ T cells were restimulated with plate-bound anti-CD3 antibodies for 24 h, and effector cytokine production was measured by ELISA.

## Discussion

Discovery of the critical role of protein S-acylation in regulation of the TCR pathway raised an important question about enzymes mediating this post-translational modification in T cells. In this study, we demonstrate that protein acyltransferase DHHC21 is an essential regulator of T cell activation that acts through S-acylation of key T cell signaling proteins – Lck, Fyn, and LAT.

Identification of several proximal TCR proteins as DHHC21 targets suggests that the lipidation machinery is a part of the TCR signaling complex and S-acylation is coordinated with other signaling events taking place immediately after TCR engagement. Indeed, we found that stimulation of the TCR prompted interaction of DHHC21 with its protein substrates and resulted in extremely rapid changes in protein S-acylation levels. This observation provides an important mechanistic detail for the previously proposed model in which S-acylation is required for compartmentalized clustering of signaling proteins in stimulated T cells (20, 33). The critical contribution of DHHC21 to TCR signaling is further supported by our data showing that loss of DHHC21 expression disrupts the initial signaling events and consequently blocks T cell activation. Furthermore, downregulation of DHHC21 removed the ability of naïve CD4^+^ T cells to differentiate into effector T helper subtypes, suggesting that this enzyme could play an important role in regulation of the T cell-mediated immunity.

Thus, this study introduces DHHC family of protein acyltransferases as a novel class of regulatory enzymes in the canonical TCR signaling pathway and establishes DHHC21 as a potential therapeutic target for diseases associated with altered T cell homeostasis.

## Experimental procedures

### Antibodies and reagents

The following antibodies were purchased from Cell Signaling: Lck (Cat. 2787), pSrc (Cat. 2101), pZAP70 (Cat. 2717), ZAP70(Cat. 3165), pErk1/2 (Cat. 4370), Erk1/2 (Cat. 4376). DHHC21 antibody was produced in our laboratory. The following reagents were purchased from Sigma-Aldrich: Protein A Sepharose (Cat. p6649), Hydroxylamine (Cat. 55459), Methyl methanethiosulfonate (MMTS) (Cat. 208795), n-Dodecyl β-D-maltoside (DDM) (Cat. D4641), Thiopropyl-Sepharose 6B (Cat. T8387), Poly-L-lysine (Cat. P8920), Phosphatase Inhibitor Cocktail 2 (Cat. P5726), cOmplete Protease Inhibitor Cocktail tablets (Cat. 11836170001). ML211 (Cat. 17630) was purchased from Cayman.

### Cells

EL-4 cells were obtained from American Type Culture Collection (Cat. TIB-39) and were grown in DMEM (Gibco, Cat. 11995–065) supplemented with 10% fetal bovine serum (Gibco, Cat. 16140–071) and glutamine (Gibco, Cat. 35050–061). EL-4 cells were maintained at densities between 0.1 and 1.0 × 10^6^ cells per ml.

For DHHC21 knockdown, a specific short hairpin RNA (shRNA) construct set was obtained from Sigma-Aldrich, and lentivirus production plasmids psPAX2 and pMD2.G were obtained from Addgene. Screening of shRNA constructs identified that clones TRCN0000121377, TRCN0000121378, TRCN0000121379, TRCN0000121380 and TRCN0000121381 resulted in a significant reduction of DHHC21 mRNA and protein levels. Clones TRCN0000121377 and TRCN0000121378 were selected to perform knockdown experiments. Lentivirus production and shRNA knockdown were performed according to protocols from Addgene (www.addgene.org/tools/protocols/plko/).

### Western blotting

Protein samples in SDS sample buffer (50 mM Tris-HCl [pH 6.8], 2% SDS, 10% glycerol, 5% β-mercaptoethanol, 0.01% bromophenol blue) were incubated at 95 °C for 5 minutes, resolved in SDS-PAGE and transferred to nitrocellulose membrane. After transfer, membranes were blocked with 5% bovine serum albumin (BSA) in PBS-T (0.1% Tween-20 in PBS buffer) and incubated with primary antibodies (1:1000) overnight, followed by three PBS-T washes. Membranes were then incubated with secondary antibodies (1:10,000) in PBS-T. After three washes in PBS-T, membranes were imaged using the LI-COR Odyssey Scanner (LI-COR Biosciences; Lincoln, NE). Brightness and contrast were adjusted in the linear range using the Image Studio software (LI-COR).

### Acyl-Resin Assisted Capture (Acyl-RAC) Assay

Protein S-acylation status was assessed by Acyl-RAC (28). Cell were lysed in 1% DDM lysis buffer (1 % Dodecyl β-D-maltoside (DDM) in DPBS; 10 μM ML211; Phosphatase Inhibitor Cocktail 2 (1:100); Protease Inhibitor Cocktail (1X), PMSF (10 mM)). Post-nuclear cell lysate was subjected to chloroform-methanol precipitation and the protein pellet was resuspended in 400 μL of blocking buffer (0.2 % S-Methyl methanethiosulfonate (MMTS) (v/v); 5 mM EDTA; 100 mM HEPES; pH 7.4) and incubated for 15 min at 42°C. MMTS was removed by three rounds of chloroform-methanol precipitation and protein pellets were dissolved in 2SB buffer (2 % SDS; 5 mM EDTA; 100 mM HEPES; pH 7.4). 1/10 of each sample was retained as input control. For thioester bond cleavage and capture of free thiol groups, freshly prepared solution of neutral 2M hydroxylamine solution (pH 7.0-7.5, final concentration 400 mM) was added together with thiopropyl sepharose to each experimental sample. In negative control samples, same concentration of sodium chloride was used instead of hydroxylamine. After 1 h incubation, beads were collected, and the proteins were eluted by incubation in SDS sample buffer supplemented with 5mM DTT and analyzed by Western blotting.

### Immunoprecipitations

Cells were lysed in 1% DDM lysis buffer (1 % Dodecyl β-D-maltoside (DDM) in DPBS; 10 μM ML211; Phosphatase Inhibitor Cocktail 2 (1:100); Protease Inhibitor Cocktail (1X), PMSF (10 mM)). 0.5 mg of protein was used for each immunoprecipitation. 1/10 of each sample was retained as input control. For immunoprecipitation, lysates were incubated with 5 μg of anti-DHHC21 antibody overnight at 4°C. Proteins were collected using 2 h incubation with protein A beads. Eluted proteins were analyzed by Western blotting.

### FACS-staining

T cell phenotype was evaluated by flow cytometry using a standardized protocol. Cells were kept on ice during all the procedures. For the extracellular markers, cells were stained with CD69 PE-Cy7 (ThermoFisher, Cat. 25-0691-82), CD25 PE (ThermoFisher, Cat. 12-0251-82). Detection of cell surface markers was conducted using a Beckman-Coulter Gallios Flow Cytometer (BD Biosciences, San Jose, CA, United States) and data were analyzed by Kaluza Analysis Software. Live/dead assays were determined using the Aqua Dead Cell Stain Kit (ThermoFisher, Cat. L34957).

### Mice

C57BL/6 mice were bred in our facility under specific pathogen-free conditions in accordance with the recommendations in the Guide for the Care and Use of Laboratory Animals of the National Institutes of Health. All the animals were handled according to approved institutional animal care protocols.

### In vitro differentiation of naïve CD4^+^ T cells

Naïve CD4+ T cells were collected from the spleen and lymph nodes of mice using naïve CD4^+^ T cell isolation kit (Miltenyi Biotech, Cat. 130-104-453) according to the instruction. Feeder cells were added to naïve T cells using splenocytes from the same strain. Feeder cells were inactivated with irradiation (30 Gy) before use. Naïve CD4+ T cells and feeder cells were mixed 1:1 with cytokine cocktails and cultured under Th1 and Th2 polarizing conditions in cell culture plates pre-coated with αCD3 (10μg/ml) and αCD28 (2μg/ml) antibodies. Th1: 50u/mL IL-2 (1μL/mL), 2ng/mL IL-12 (2μL/mL), 10μg/mL αIL-4 (10μL/mL), 10ng/mL IFN-γ (1μL/mL) Th2: 50u/mL IL-2 (1μL/mL), 10ng/mL IL-4 (1μL/mL), 10μg/mL αIFN-γ (10μL/mL), 20ng/mL IL-6 (2μL/mL). After 4 days cells were re-stimulated and cytokine production was measured by ELISA (R&D Systems).

### Fura-2 Calcium Imaging

Cells were loaded with Fura-2 AM as described previously (34, 35) and placed in an imaging chamber with a glass bottom pre-coated with Poly-L-lysine. Images were taken on a Nikon TiS inverted microscope (Tokyo, Japan) with a 40× oil immersion objective, and images were taken every second with a Photometrics Evolve electron-multiplying charged-coupled device camera (Tucson, AZ). Cells were pre-treated with αCD3 antibody and secondary IgG antibody was added after baseline recording to initiate TCR signaling.

## Acknowledgements

We would like to thank Savannah West for critical reading of the manuscript.

## FOOTNOTES

This work was supported by National Institute of General Medical Sciences grants 1R01GM115446-01A1 (to A.M.A.), 1R01GM130840 (to A.M.A., D.B.), and 2R01GM081685 (to D.B.).

The abbreviations used are: TCR, T cell receptor; LAT, linker for activation of T cells; PAT, protein acyltransferase; Th, T helper.

