## Supplementary Figures S1 and S2 for "Regulation of T cell receptor signaling by protein acyltransferase DHHC21"

**Supporting information**

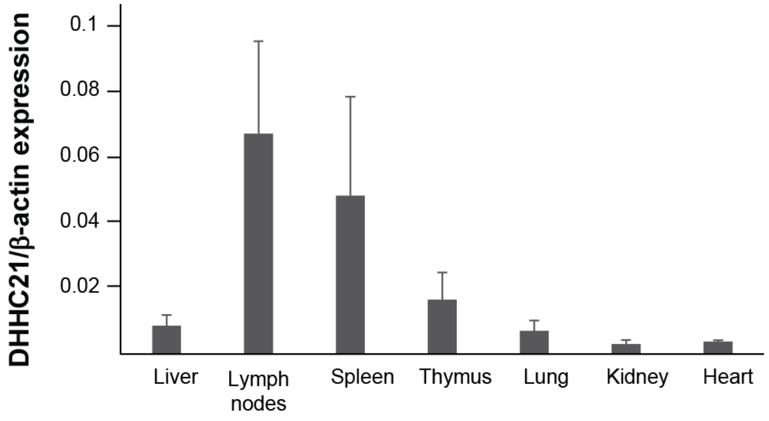


**Figure S1.** DHHC21 mRNA expression profile in murine tissues, shown relative to β-actin.

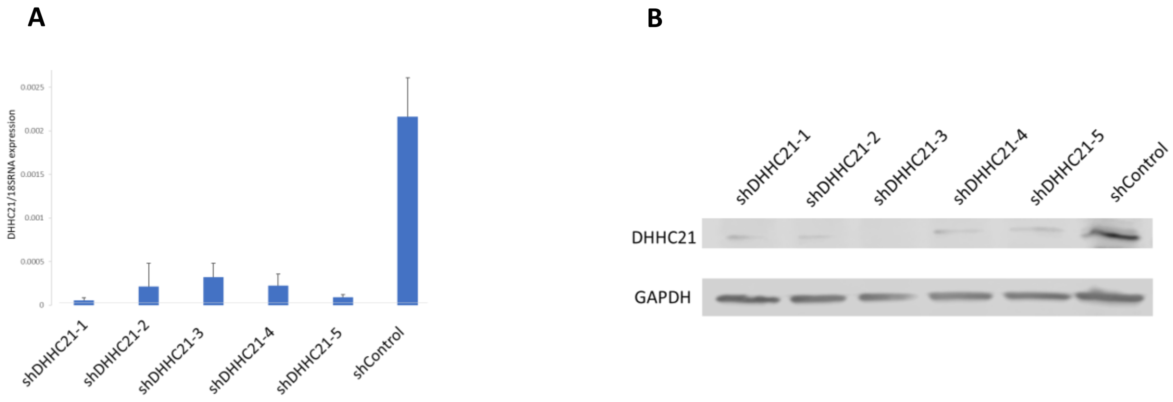


**Figure S2. shRNA-mediated knockdown of DHHC21.** Murine EL-4 T cells were transduced by lentivirus carrying DHHC21-targeting or control nonspecific shRNA. Downregulation of DHHC21 expression in EL-4 T cells was confirmed by **(A)** qPCR analysis of mRNA levels and **(B)** Western blotting.
